# Large portion of essential genes is missed by screening either fly or beetle indicating unexpected diversity of insect gene function

**DOI:** 10.1101/2021.02.03.429118

**Authors:** Muhammad Salim Hakeemi, Salim Ansari, Matthias Teuscher, Matthias Weißkopf, Daniela Großmann, Tobias Kessel, Jürgen Dönitz, Janna Siemanowski, Xuebin Wan, Dorothea Schultheis, Manfred Frasch, Siegfried Roth, Michael Schoppmeier, Martin Klingler, Gregor Bucher

**Author notes:** (equally contributing) (Muhammad Salim Hakeemi, Salim Ansari, Matthias Teuscher, Matthias Weißkopf).

## Abstract

Most gene functions were detected by screens in very few model organisms but it has remained unclear how comprehensive these data are. Here, we expanded our RNAi screen in the red flour beetle *Tribolium castaneum* to cover more than half of the protein-coding genes and we compared the gene sets involved in several processes between beetle and fly.

We find that around 50 % of the gene functions are detected in both species while the rest was found only in fly (^~^10%) or beetle (^~^40%) reflecting both technical and biological differences. We conclude that work in complementary model systems is required to gain a comprehensive picture on gene functions documented by the annotation of novel GO terms for 96 genes studied here. The RNAi screening resources developed in this project, the expanding transgenic tool-kit and our large-scale functional data make *T. castaneum* an excellent model system in that endeavor.

## Introduction

Only in a very small number of genetic model species like the mouse *Mus musculus*, the zebrafish *Danio rerio*, the nematode *Caenorhabditis elegans* and the vinegar fly *Drosophila melanogaster* have the functions of most genes been assayed in systematic screens. This restriction to few model systems is a consequence of the necessity for an elaborate genetic and molecular tool kit, which is extremely laborious to establish (Jorgensen and Mango, 2002; Kile and Hilton, 2005; Patton and Zon, 2001; St Johnston, 2002). Unfortunately, it has remained unclear how representative findings in these model species actually are for their clade or in other words, how quickly and profoundly gene function diverges in evolution. Knowing the degrees of gene function divergence is relevant not only for understanding the evolution of biodiversity but also for applied research, e.g. for transferring knowledge from model systems to species relevant for medical applications or pest control.

Recently, the study of gene function has been extended to non-traditional model organisms. Predominantly, candidate genes known for their function in the classical model systems have been tested in other organisms. Subsequent comparisons revealed both, conservation and divergence of gene functions. For example, axis formation in *D. melanogaster* has turned out to be a rather diverged process partially based on different genes compared to other insects. The key anterior morphogen of *D. melanogaster, bicoid*, is not present in most insects (Brown et al., 2001). Instead, repression of Wnt signaling plays a central role in the red flour beetle *Tribolium castaneum* (Fu et al., 2012) as it does in many animals including other insects, flatworms and vertebrates (Glinka et al., 1998; Gurley et al., 2008; Klomp et al., 2015; Yoon et al., 2019) - but not in *D. melanogaster*. The functions of genes of the Hox cluster, in contrast, appear conserved over very large phylogenetic distances - although some functional divergence has been linked to the evolution of arthropod morphology (Averof, 2002). Likewise, the gene regulatory network of dorso-ventral patterning and head specification show the involvement of similar gene sets, although a few components appear to be involved in only some clades (Kittelmann et al., 2013; Kitzmann et al., 2017; Lynch and Roth, 2011; Stappert et al., 2016).

Notably, the differences in gene functions documented so far may be an underestimation of the real divergence, because the prevailing candidate gene approach leads to a systematic bias towards conservation. The genes to be tested are usually chosen based on the knowledge of their ortholog’s involvement in other species. As a consequence, unrelated genes are rarely tested and the involvement of unexpected genes in a given process is underestimated. Hence, approaches are needed to overcome this bias and to gain a realistic view on the degree of gene function divergence. To that end, genes required for certain biological processes need to be identified in an unbiased and genome-wide manner also in non-traditional organisms, even though this has remained technically challenging.

The red flour beetle *T. castaneum* has recently been established as the only arthropod model organism apart from *D. melanogaster* where genome-wide unbiased RNAi screens are feasible. Based on the robust and systemic RNAi response of this species, the *iBeetle* large scale screen was performed where random genes were knocked down and the resulting animals were scored for a number of developmental phenotypes (Bucher et al., 2002; Schmitt-Engel et al., 2015; Tomoyasu and Denell, 2004). Apart from its particularly strong and robust RNAi response, *T. castaneum* offers a comparably large tool kit for analyzing gene function including transgenic and genome editing approaches (Berghammer et al., 1999; Gilles et al., 2015; Schinko et al., 2010).

In this paper, we used an expanded dataset to assess the degree of divergence of the gene sets involved in selected developmental processes between fly and beetle such as head, muscle and ovary development, and dorso-ventral patterning. First, we determined genes that were essential in the beetle for these processes but which had so far not been connected to them in *D. melanogaster*. These a priori unexpected genes sum up to about 37% of the total genes identified to be involved in either one or both species. For 30% of these genes, no functional annotation had been available at FlyBase at all such that we provide the first functional Gene Ontology (GO) assignment for the respective ortholog group in insects. Only two genes essential in *T. castaneum* did not have an ortholog in *D. melanogaster*, i.e. these processes seem not much affectd by gene gain or loss. We conclude that restricting genetic screens to one model system only, falls short of identifying a comprehensive set of essential genes. Further, our data reveals an unexpected degree of divergence of gene function between two holometabolous insect species. We also present here an update of the dataset gained in the genome wide iBeetle screen in *T. castaneum*. Our analysis is based on both, a dataset previously published comprising 5.300 genes (Schmitt-Engel et al., 2015) and an additional 3.200 genes screened as part of this project. In addition to those, we also make accessible (at iBeetle-Base) the phenotypes for an additional 4,520 genes which were screened while the analysis presented here was ongoing. Hence, with this paper, the coverage of genes tested and annotated at iBeetle-Base sums up to 13.020 Tribolium genes (78 % of the predicted gene set).

## Results

### Continuation of the large scale iBeetle screen

We added 3,200 genes to the previously published 5,300 genes of our large scale *iBeetle* screen (Schmitt-Engel et al., 2015), reaching a coverage of 51% of the *T. castaneum* gene set of total 16,593 currently annotated genes (Herndon et al., 2020). We followed the previously described procedure for the pupal injection screen (Schmitt-Engel et al., 2015) with minor modifications (see methods). In short, we injected 10 female pupae per gene with dsRNAs (concentration 1ug/ul). We annotated the phenotypes of the injected animals and the first instar cuticle of their offspring using the EQM system (Mungall et al., 2010), the *T. castaneum* morphological ontology *Tron* (Dönitz et al., 2013) and a controlled vocabulary (see Schmitt-Engel et al. 2015). The data is available at the online database *iBeetle-Base* (http://ibeetle-base.uni-goettingen.de/) (Dönitz et al., 2015; Dönitz et al., 2018). Our controls revealed a similar portion of false negative and false positive annotations as in the first part of the screen (Fig. 1 and Table S1). The detailed analysis presented below was based on this set of genes covering approximately 50% of the genome. In parallel, we continued the screen and have in the meanwhile reached a coverage of 78 % (13,020 genes). We publish these additional phenotypic data (accessible online at *iBeetle-Base*) with this article, but they were not included in the detailed analysis presented here because both analyses ran in parallel.

**Figure 1:**
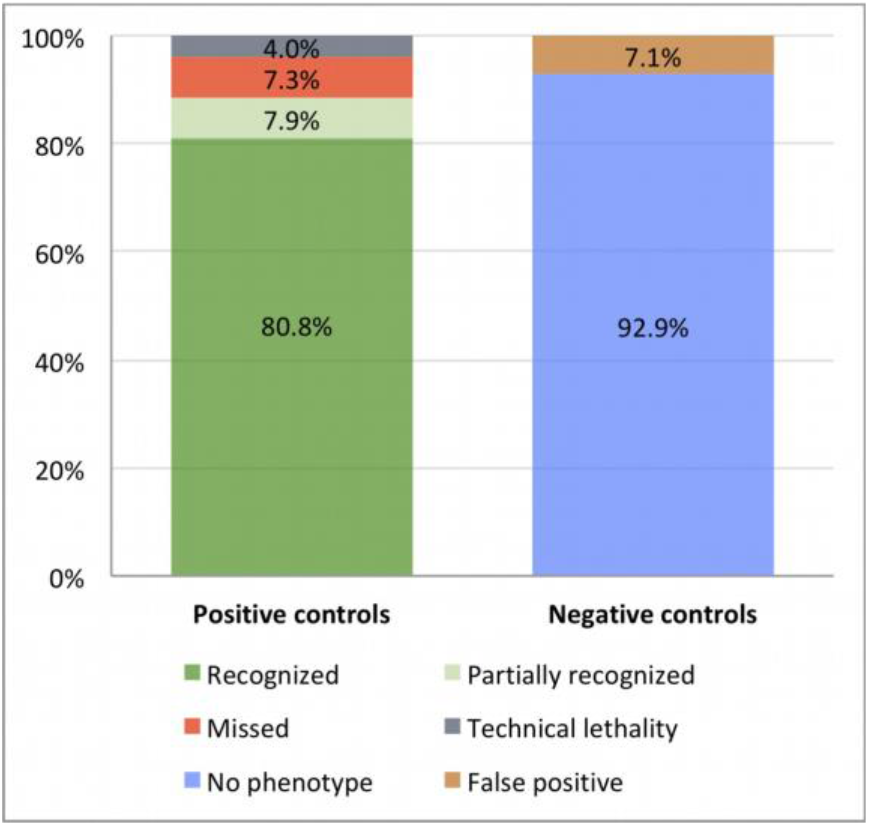
Quality controls of the primary screen. 178 positive controls using 35 different genes were included. More than 88% of the positive controls were fully or partially recognized (left bar) while 7.3% were missed. 4% could not be analyzed due to technical lethality before the production of offspring. 7.1% of the negative controls were false positively annotated (right bar). These figures are similar to the first screening phase (Schmitt-Engel et al., 2015).

### Unexpected gene functions in developmental processes

We wanted to use our large-scale phenotypic dataset to systematically compare the gene sets involved in the same biological processes in *T. castaneum* and *D. melanogaster*. To that end, we first identified in an unbiased way all genes involved in a number of biological processes by searching *iBeetle-Base*. Specifically, we scored for phenotypes indicative of functions in dorso-ventral patterning, head and muscle development, in oogenesis, and epithelial adhesion in wings (wing blister phenotypes). For all these processes, we found a number of gene functions that were expected based on *D. melanogaster* knowledge (see below). This confirmed that the screen design allowed detection of respective phenotypes. Importantly, we also found functions for genes so far not connected to those processes (based on FlyBase information, PubMed searches and scientist expertise). The *iBeetle* screen is a first pass screen with a focus on minimizing false negative results with the trade-off of allowing for false positive annotations (Schmitt-Engel et al., 2015). The likelihood for this type of error is further increased by off-target effects and/or by strain specific differences in the phenotype (Kitzmann et al., 2013). Hence, we aimed at excluding false positive annotations for the unexpected gene functions. First, we based our analyses only on genes for which phenotypes had been annotated with a penetrance of >50% in the primary screen. Further, we only used phenotypes that were reproduced by RNAi experiments with non-overlapping dsRNA fragments targeting the same gene. In order to exclude genetic background effects, we used another lab strain (our standard lab strain *San Bernardino, SB*) except for the muscle project where we needed to use the *pBA19* strain, which has EGFP marked muscles (Lorenzen et al., 2007). This re-screening procedure resulted in a set of genes for which we can claim with high confidence that they are indeed involved in these processes in *T. castaneum* - but which previously were not assigned to these in *D. melanogaster* (Supplementary Table S2).

### Assigning the first function to a gene versus extending previous annotations

One reason for a lack of respective functional data in FlyBase could be that the knocked-down beetle gene does not have an ortholog in the fly. In order to test this hypothesis, we searched for the fly orthologs in orthoDB and by manually generating phylogenetic trees based on searching *T. castaneum, D. melanogaster* and *M. musculus* genomes for orthologs and paralogs. This analysis revealed that only three genes with a novel function (appr. 3%) did not have a *D. melanogaster* ortholog (yellow in Fig. 2). Evidently, lineage-specific gene loss or gain explains only a minor part of the functional divergence of homologous developmental processes.

**Figure 2.**
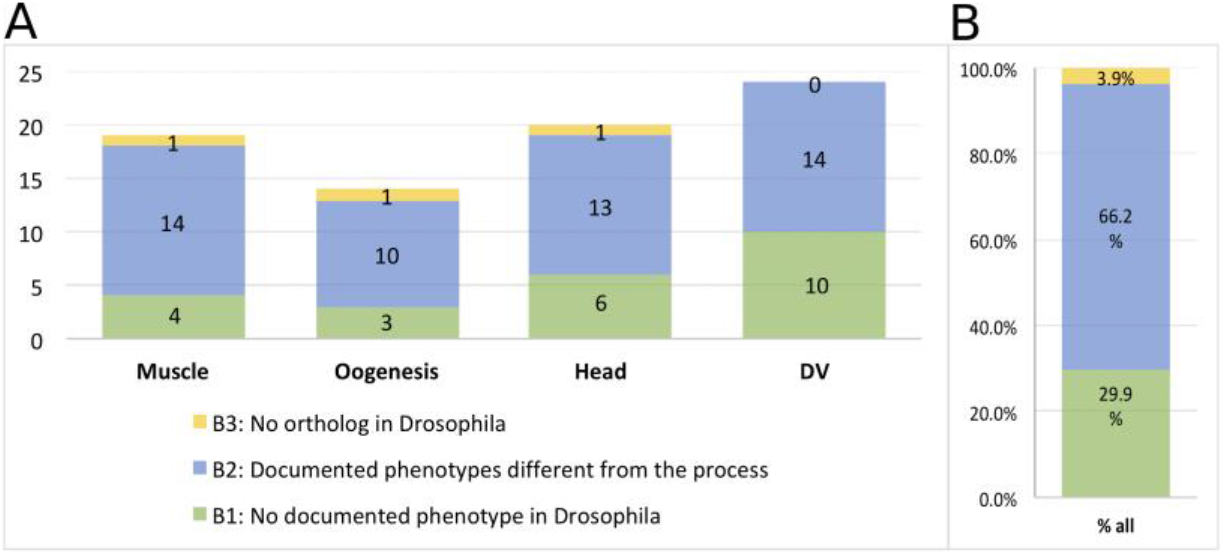
Analysis of genes with unexpected gene functions. A) Numbers of genes with unexpected function in the respective process. B) Combined numbers for all four processes. Only a small portion of genes with novel gene functions did not have orthologs in *Drosophila* (yellow). About two-thirds of the genes had previous phenotypic annotations relating to other biological processes (blue). For one third of those genes, we had detected the first phenotype for this gene within insects (green).

Next, we asked whether the respective *D. melanogaster* orthologs were known to be involved in other biological processes or lacked any phenotype information. To that end, we looked up phenotype information of the respective *D. melanogaster* orthologs on FlyBase (analysis done with OrthoDB v9). Among the fly orthologs whose functional annotations did not match with those from the iBeetle screen or published record, around two thirds (64.6 %) had annotations that were related to other processes than the ones studied in *T. castaneum* (Fig. 2). Importantly, one third of the genes (32.3 %) did not have any functional annotation in FlyBase. Hence, for those genes, the *iBeetle-screen* had detected the first documented function of that ortholog group in insects. Importantly, due to the lack of previous phenotypic information, these genes likely would not have been included in a classical candidate gene approach.

### A quarter of *Drosophila* gene function annotations were not confirmed for *T. castaneum*

In a complementary approach, we asked how many genes known to be involved in a given process in *D. melanogaster* had been assigned related functions in the *iBeetle* screen. To that end, we first collected lists of genes involved in those processes based on *D. melanogaster* knowledge (expert knowledge, literature and FlyBase) (Table S3). Then we mined *iBeetle-Base* to see how many of the beetle orthologs had an annotation related to that process (Fig. 3A). About two-thirds of those genes had actually been screened in *T. castaneum* (Fig. S1) and all following numbers are based on the analysis of this subset.

**Figure 3.**
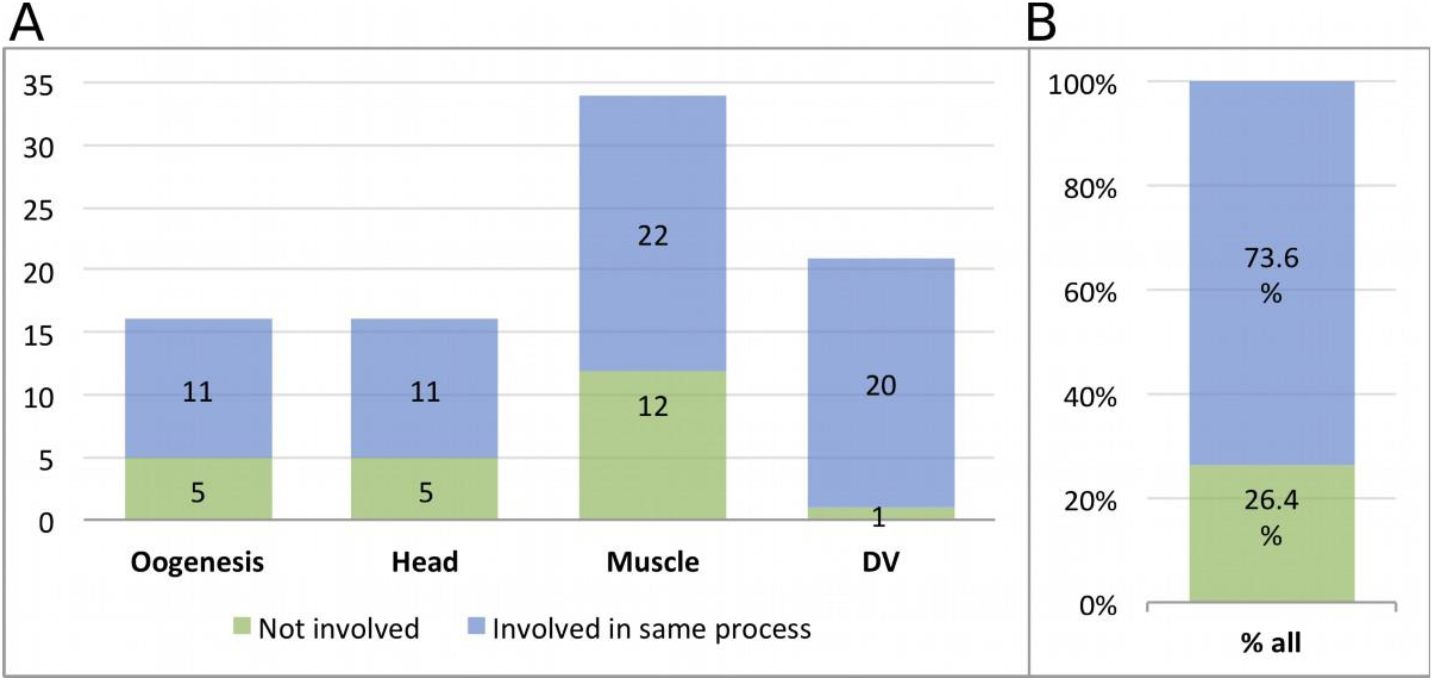
Beetle genes showing phenotypes expected from *Drosophila*. A) Gene sets known to be involved in given processes in *Drosophila* were compared to iBeetle data. Many showed related phenotypes (blue) while others had no or different types of phenotypes (green). B) Approximately one quarter of the genes known to be involved in certain *Drosophila* processes were not required in that process in *Tribolium*. This analysis is based on the subset of genes which already had been screened in *Tribolium* (66%).

A surprisingly large portion of genes (26.4%) known to be involved in these processes in *D. melanogaster* did not show the expected phenotype in *T. castaneum* (Figure 3B).

### Enriching the GO information with data from *Tribolium*

Gene ontology (GO) assignment is a powerful tool to establish hypotheses on the function of given gene sets (Carbon et al., 2009). So far, there were no GO terms associated based on *T. castaneum* data. The work presented here revealed that a surprisingly high portion of orthologous genes has diverging functions in different organisms. To enrich the GO database, we submitted GO terms with respect to the biological process for all 96 re-screened genes with functions in dorso-ventral patterning (GO:0010084), oogenesis (GO:0048477), the development of embryonic muscles (GO:0060538) and head (GO:0048568).

[[the new GO terms are submitted but not yet accepted. This part will only be included in the final version of the paper if the terms have been accepted by the GO consortium]]

## Materials and Methods

### Screen

We followed the tested and published procedures apart from some minor changes (please find an extensive description of the procedure in Schmitt-Engel et al. 2015). In particular, we used the same strains, injection procedures, and incubation temperatures and incubation times. dsRNAs were produced by Eupheria Biotech Dresden, Germany. Different from the published procedure, the stink gland analysis was performed 21 days after pupal injection (in the first screening phase, this analysis had been performed after larval injection).

### Controls of the screen

To assess the sensitivity and reliability of the screen, and also to test the accuracy of each screener, we included approximately 5% positive controls from a set of 35 different genes. By and large, we used the same positive controls as in the first screening phase (see Table Table_S1_controls). *Tc-zen-1* was excluded since the phenotypes were much weaker than in the previous screen, probably due to degradation of the dsRNA. We added new positive controls to score for muscle and stink-gland phenotypes, which we took from novel genes detected in the first screening phase. Muscle phenotypes iB_06061, iB_05796, iB_03227, iB_01705; stink gland and ovary phenotypes: iB_02517. Head defects: iB_05442 (that gene was not scored for its stink gland phenotype because it turned out to be too mild to be identified reliably in high throughput). In 143 cases (80.8%, n=177), the phenotypes of positive controls were fully recognized (for comparison: in the first screening phase the respective numbers were: 90%, n=201). In 14 cases (7.9%; phase 1: 4%) the phenotype was partially recognized. This category includes complex phenotypes where half (one of two aspects: *knirps, piwi, SCR, cta, cnc, iB_01705, iB_05442*) or two of three aspects (*aristaless*) of all phenotypic aspects were correctly identified. 13 phenotypes were missed completely (7.3%, phase 1: 4%). *Tc-metoprene tolerant (Tc-met*) was missed most frequently, probably due to the fact that the embryonic leg phenotype was very subtle and in addition, the penetrance of the phenotype appeared to be lower than in the first screen (penetrance: less than 30%). Seven positive controls (4%, phase 1: 1%) could not be analyzed due to prior technical lethality, i.e. the premature death of the injected pupae prevented the detection of the phenotype. In three cases wrong aspects were annotated (false positive: 1.7%). Depending on the other annotations these positive controls were valued as partially recognized (SCR) or missed (met, CTA). Find more details in Table Table_S1_controls.

Negative controls (buffer injections) were mainly annotated correctly (no phenotype in 92.9%; phase 1: 96%) and just in 7 cases led to false positive annotations (7.1%; phase 1: 2%) (Table Table_S1_controls; sheet 2).

### Re-Screen

Re-screening of selected iBeetle candidates involved in a number of biological processes was performed in order to probe for off-target and strain-specific effects. For that purpose, two independent dsRNA fragments (original iB-fragments and one non-overlapping fragment, both at concentration 1 μg/μl) of the same gene were injected separately into a different genetic background (*San Bernardino, SB* strain), except for the muscle project where it is required to use the pBA19 strain with EGFP marked muscles. The rest of the injection procedures and analyses were as in the first pupal injection experiment (see details in Materials and Methods).

### Phylogenetic analysis

The *Tribolium* protein sequences from gene set (http://ibeetle-base.uni-goettingen.de/downloads/OGS3_proteins.fasta.gz-includingchangesfrom2016/02/15) were used to retrieve the most similar proteins of *T. castaneum, D. melanogaster* and *M. musculus* excluding isoforms. Multiple alignments were done with the ClustalOmega plugin as implemented in the Geneious 10.1.3 software (Biomatters, Auckland, New Zealand) using standard settings. Alignments were trimmed to remove poorly aligned sequence stretches. Phylogenetic trees were calculated using the FastTree 2.1.5 plugin implemented in Geneious.

### Generation of Unc-76 mutations via CRISPR/Cas9

The procedure used to generate *Unc-76* mutations was described by Basset et al., 2013(Bassett et al., 2013). For making the template for the guide RNAs, the *Unc-76* target sequence between the T7 promoter and the gRNA core sequence in the forward primer, gRNA_F, was chosen as GGTTCAACGATCTGACCAGTG, and after annealing gRNA_F with SGRNAR the template was PCR amplified with Q5 polymerase (NEB). Guide RNAs were transcribed with Ampliscribe T7 Flash (epicentre), isolated with the MEGAclear kit (Ambion), and injected together with Cas9 mRNA into *w*[1118] *sn*[3] *P{ry+t7.2=neoFRT}19A* embryos. Single lines established from the offspring were tested as heterozygotes over *FM7c* with the T7 endonuclease assay for sequence alterations near the target site (Kondo and Ueda, 2013). The lethal *Unc-76*[CR007] allele carries a 16 nucleotide deletion near the target site in the sequence ..TAT CCA CAC ACc aac ggt ttg gga tcc GGA TCC GGA TCC.. of the second exon (X: 2091152... 2091167, r6.32; see lower case letters) that creates a frameshift in the ORF in all known isoforms (after T246 in Unc-76 RA to -C and after T61 in Unc-76 RD).

## Discussion

### Investigating one species falls short of a comprehensive view on gene function

Large scale screens in the leading insect model organism *D. melanogaster* have revealed gene sets involved in certain biological processes. As consequence, insect-related GO term annotations are almost exclusively based on work in flies. However, there are several reasons to believe that the picture has remained incomplete. On one hand, species-specific or technical limitations may have prohibited identification of an involved gene in *D. melanogaster*. On the other hand, evolution may have led to functional changes such as the loss of ancestral gene functions or the integration of genes into a novel process. Unfortunately, it has remained unclear to what extent the gene sets determined exclusively in flies would be representative of insects as a whole.

Our systematic screening in a complementary model organism has revealed that the identified gene sets show an astonishing degree of divergence. Based on our calculations (see details below) we estimate that only half of the gene functions are similarly detected in both species (52%, column 4 of Fig. 4A) while the remaining gene functions were revealed either only in *D. melanogaster* (11%, column 4 of Fig. 4A) or only in *T. castaneum* (37%, column 4 of Fig. 4A). Hence, our current knowledge based on screening in one species appears to be much less comprehensive than previously thought. We believe that the different proportions of genes detected in only one species (11% vs. 37%) may reflect both, biological and technical differences (see below).

**Figure 4.**
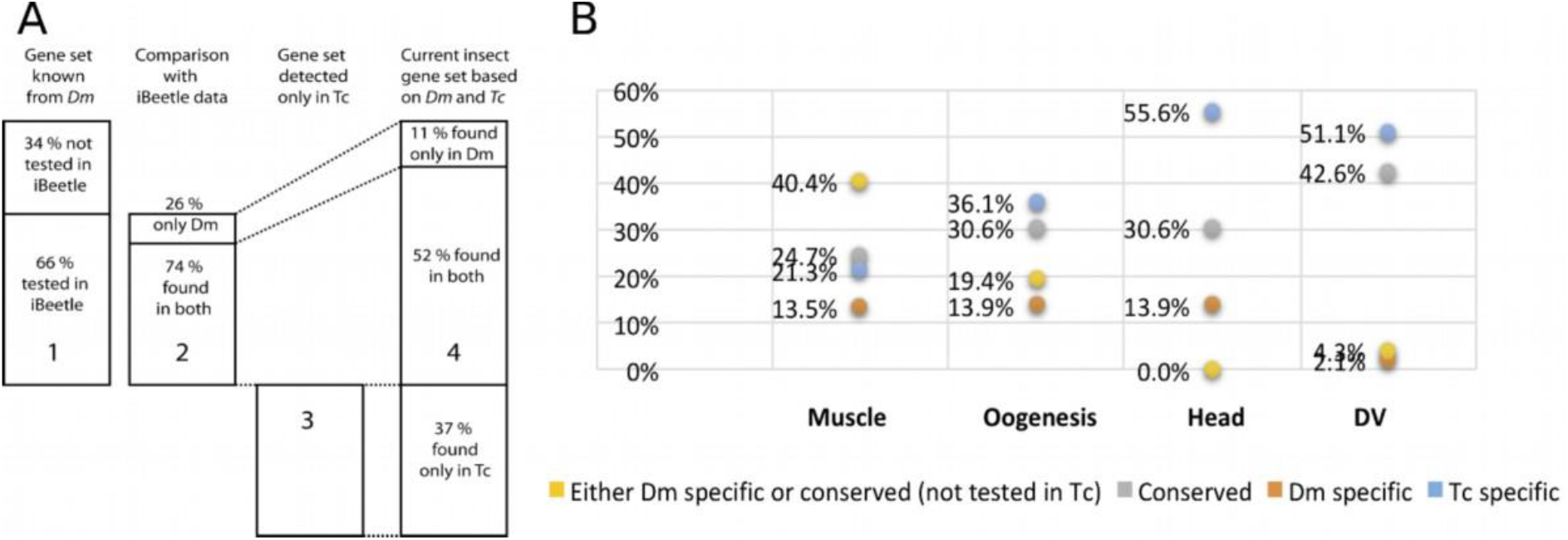
Many genes are detected only in one of the species in the same processes. Combining genes found in fly (column 1) and/or beetle (column 3) leads to the currently known insect gene set for the processes analysed here. Portions shown in column 1 and 2 are based on Fig. S1, Fig.2 and Fig. 3. We calculate the portions of genes of the combined insect gene set (column 4), which were detected only in *Drosophila* (11 %), only in *Tribolium* (37%) or in both (52%). See text for details and discussion of potential systematic biases. B) Respective values for the single processes show that the minimum contribution of the *Tribolium* screening platform amounted to 20% genes not detected in *Drosophila*. See table S4 for calculations. Neither model species is able by itself to detect “the insect gene set”.

In summary, despite some uncertainties with respect to the exact numbers (see discussion below), our findings provide a compelling argument that focusing on single model species falls short of comprehensively revealing the genetic basis of biological processes in any given clade. Further, it shows that *T. castaneum* is an extremely useful screening system for insect biology, able to reveal novel gene functions even in processes that have been studied intensely in *D. melanogaster*.

### Estimating the portions of gene functions revealed in fly versus beetle

Our beetle data are based on both, our systematic screening of 51% of the *T. castaneum* gene set and on previous candidate gene work. With respect to fly data, we rely on information available on FlyBase and our expert knowledge of the processes under scrutiny. Given these different kinds of sources and approaches, the data are prone to various types of uncertainties. Therefore, we discuss the way we combined the numbers to calculate our estimation. Subsequently, we will discuss some uncertainties and in how far they influence the estimation.

Of the genes known from *D. melanogaster* to be involved in the processes investigated here (n = 132; see Table S4), we could compare 66% to iBeetle data (column 1 in Fig. 4A; based on Fig. S1; n = 87). Of those genes, 26% (n = 23) were not involved in that process in *T. castaneum* (column 2 in Fig. 4A; based on Fig. 3). For our overall estimation, we extrapolated this share to the total number of genes involved in the fly (hatched lines from column 2 to column 4). A number of gene functions detected in the iBeetle screen had not been assigned such functions in *D. melanogaster* before (column 3 in Fig. 4A; based on Fig. 2). When combining these numbers, we aimed at providing a minimum estimation for divergence of detected gene functions (Column 4 in Fig. 4A). To be conservative, we assumed that all gene functions known from *D. melanogaster* but not yet tested in the iBeetle screen would fall into the class of genes being involved in both species (see numbers in green square in Table S4). Further, we scored each signaling pathways as one case (finding mostly conservation) even if single components of these pathways had not divergent phenotypes. This conservative assumption leads to the abovementioned minimum estimation of divergence in these gene sets (Column 4 in Fig. 4A; calculation in Table S4). Of all genes currently known to be involved in one of the processes we studied, the portion of genes detected exclusively in the fly (11%; n = 23) is much smaller than the one detected only in the beetle (37%; n = 76) while the analogous function of half of the genes (52%; n = 109) is detected in both species.

With this work, we present the first and a quite extensive dataset to estimate this kind of numbers. Still, some confounding issues need to be considered. The first uncertainty stems from the fact that the beetle data is based on testing about 50 % of the genes. In the second part of the screen, we had prioritized genes that were e.g. highly expressed, showed sequence conservation and had GO annotations. The prioritization apparently was successful as 66% of the gene functions known from *D. melanogaster* had been covered in the iBeetle screen (Fig. 4A), which is much more than the 40% expected for an unbiased selection (Schmitt-Engel et al., 2015). Hence, our figures might be biased towards conserved gene function. As a consequence, the overall portion of beetle specific genes without conserved functions likely is even higher than reflected in Fig. 4A.

Second, we found quite different numbers for the four processes under scrutiny (Fig. 4B). However, even in the process with the lowest portion of genes detected exclusively in *T. castaneum* (muscle development), this portion was 21%, which still indicates a significant degree of unexplored biology.

Third, the *D. melanogaster* numbers could be influenced by false negative data. The data on FlyBase has not been gathered in one or few standardized screens where all data were published – it is mainly based on published results of single gene analyses. However, not all genetic screens have reached saturation and not all genes detected in large-scale screens may have been further analyzed and published. Hence, the number of genes in principle detectable in *D. melanogaster* might actually be larger than the numbers extracted from FlyBase. In the iBeetle screen, in contrast, negative data was systematically documented, such that this type of uncertainty is restricted to technical false negative data, which we found to be around 15% in this first pass screen (Fig. 1). This uncertainty could potentially increase the portion of *D. melanogaster* specific or conserved genes. Fourth, theoretically there may be false positive data albeit restricted to the set of genes detected in both species. The reason is that iBeetle was a first pass screen, where we aimed at reducing false negative data with the tradeoff that false positive data are enriched (Schmitt-Engel et al., 2015). Although finding similar phenotypes in two different species will not in many cases be false positive, we tried to minimize this error by manually checking the annotations of the respective genes, excluding those that showed a phenotype with low penetrance or in combination with many other defects indicating a non-specific effect. Of note, the issue of false positives is restricted to the genes detected in both species (column 2; based on Fig. 3). It does not apply to those genes detected only in the beetle but not the fly (column 3; based on Fig. 2) because in this case, all phenotypes were confirmed by independent experiments with non-overlapping dsRNA fragments in different genetic backgrounds such that false positive results are excluded. In summary, while there are a number of uncertainties that we could not clarify with available data or methods, most of these uncertainties hint at underestimation rather than overestimation of functional divergence between fly and beetle.

### Technical characteristics contribute to the detection of unequal gene sets

Our numbers reveal that functionally comparable gene sets in two quite closely related model systems are far from identical. A question of obvious biological relevance but not easily resolved is: to which degree do these differences reflect biologically meaningful divergence of gene functions, or alternatively, simply result from technical problems, i.e. reflect different strengths and weaknesses of the respective screening methods and model systems?

As discussed above, some degree of false negative data may be expected in both model systems. In case of the iBeetle screen, this will be restricted to technical false negative data. In the *D. melanogaster* field, there may be additional false negative data due to the lack of saturation of screens and/or lack of reporting of genes that were not studied in detail. However, given the extent and comprehensiveness of work in the *D. melanogaster* field we feel that this might not be of high relevance. As to different strengths of screening procedures, it is certainly true that the way screens are performed influences what sets of genes can be detected. For instance, our parental RNAi approach knocked down both, maternal and zygotic contributions while some classic *D. melanogaster* screens affected only the zygotic contribution. Hence, genes where maternal contribution rescues the embryonic phenotype are easily missed in the fly but not the beetle. For instance, parental RNAi knocking down components of the aPKC complex leads to severe early disruption of embryogenesis in *T. castaneum* while in respective *D. melanogaster* mutants almost no defects are seen on the cuticle level (A. Wodarz, unpublished observation). Conversely, our RNAi screen depended on the accuracy of gene annotations and our approach of screening for several processes in parallel may have reduced detection sensitivity. One striking example for the different strengths of screening designs is provided by wing blister phenotypes. In the first part of the *iBeetle* screen we detected 34 genes showing wing blister phenotypes where 14 did not have related GO term annotation at FlyBase and 5 did not have any GO annotation at all. Seven of these genes were subsequently tested by RNAi lines in *D. melanogaster* where four of them indeed showed a related phenotype. Likewise, some wing blister genes from *D. melanogaster* were not annotated in the iBeetle screen. When we checked more specifically, this was often due to lethality of the animal before the formation of wings (Schmitt-Engel et al., 2015). When we varied the timing of injection, two of those knock-downs elicited wing blister phenotypes also in *T. castaneum* (Schmitt-Engel et al., 2015). These data show that details of the screening procedure influence the subset of genes that is detected.

### Evolutionary divergence of gene function and derivededness of *Drosophila* biology may be larger than appreciated

Most relevant for the field of functional genetics is our conclusion that the degree of divergence of gene functions is larger than previously assumed. Therefore, some genes are detected only in one species because the gene’s function is not required for that process in the other. Indeed, there is evidence supporting this view. In a recent study, a number of muscle genes identified in the *iBeetle* screen were more closely investigated in *D. melanogaster* (Schultheis et al., 2019a; Schultheis et al., 2019b). Despite some efforts, the negative data for fly orthologs appeared to be real negative. For example, null mutations of one of the genes found in our beetle, *nostrin*, did not elicit a phenotype in *D. melanogaster* unless combined with a mutation of a related F-bar protein *Cip4*. Likewise, *Rbm24* displays strong RNAi and mutant phenotypes in *T. castaneum* and vertebrates, respectively, but *D. melanogaster* is lacking an *Rbm24* ortholog, and functional compensation by paralogs was suggested to occur during *D melanogaster* muscle development. Other genes including *kahuli* and *unc-76* are expressed in the *D. melanogaster* mesoderm but only showed very subtle somatic muscle phenotypes, if any, in Mef2-GAL4 driven RNAi experiments or with CRISPR/Cas9 induced mutations, respectively (see Materials & Methods). By contrast, their beetle counterparts had strong and penetrant phenotypes in single knock-downs (Schultheis, 2016; Schultheis et al., 2019a; Schultheis et al., 2019b). These data suggest that the function of genes or their relative contribution to this biological process have changed significantly. They also indicate that the single gene view may be limited. Phenotypes depend on networks of interacting genes and this may allow for changes and replacements of individual components while the overall network structure is maintained. There are more striking examples of gene function changes. The gene *germ cell-less* was detected in the iBeetle screen to govern anterior-posterior axis formation in the beetle while in *D. melanogaster* it is required for the formation of the posterior germ-cells (Ansari et al., 2018). Also, the *D. melanogaster* textbook example of a developmental morphogen *bicoid* does not even exist in *T. castaneum* (Brown et al., 2001) and yet other genes were found to act as anterior determinants in other flies (Klomp et al., 2015; Yoon et al., 2019). Along the same lines, the genes *forkhead* and *buttonhead* do not appear to be required for anterior patterning in *T. castaneum* but are essential in flies (Kittelmann et al., 2013; Schinko et al., 2008; Weigel et al., 1989; Wimmer et al., 1997).

These findings with respect to specific genes add to a number of observations arguing for a comparatively high degree of derivededness of fly biology. The number of genes is much smaller in *D. melanogaster* (appr. 14,000) compared to *T.castaneum* (appr. 16,500). Further, a number of developmental processes are represented in a more insect-typical way in *T. castaneum* like for instance segmentation (Tautz et al., 1994), head (Posnien et al., 2010) and leg development, brain development (Farnworth et al., 2019), extraembryonic tissue movements (Panfilio, 2008) and mode of metamorphosis (Snodgrass, 1954). In most cases, the situation in the fly is simplified and streamlined for faster development.

We think that these biological difference lead to divergence in gene function, which we just started to uncover. Given the large divergence of gene sets found in different screening systems, and the documented cases of biological divergence of gene function, we propose that a more systematic investigation on the divergence of gene function is needed and that hypothesis independent screening now possible in *T. castaneum* may be helpful in that endeavor.

## Supporting information

Table_S1_controls_v1

Table_S2_Comparison Tribolium Phenotype with Drosophila data

Table_S3_Drosophila genes compared to Tribolium_iBeetle_annotation

Table_S4_combined analysis

## Author contributions

Data presentation and writing of manuscript:

*Muhammad Salim Hakeemi, G.B*.

Screen and Re-Screen; analysis of entire set of candidates:

*Muhammad Salim Hakeemi, Salim Ansari, Matthias Teuscher, Matthias Weißkopf*

Data handling and processing

*Jürgen Dönitz*

Data collection and analysis for Drosophila comparison:

*Janna Siemanowski, Daniela Großmann, Muhammad Salim Hakeemi*

*Annotation as part of Bayer screen*

*Xuebin Wan*

Generation of Drosophila mutant

*Dorothea Schultheis*

Supervision and interpretation of Re-Screen:

*Martin Klingler, Michael Schoppmeier, Gregor Bucher*

*Siegfried Roth, Manfred Frasch*

Supervision primary Screen:

*Daniela Großmann, Tobias Kessel (geb. Richter)*, Michael Schoppmeier, Martin Klingler, Gregor Bucher

Coordination of screen funding and realization

Martin Klingler, Michael Schoppmeier, Gregor Bucher

## Author contributions

Data presentation and writing of manuscript:

*Muhammad Salim Hakeemi, Gregor Bucher*

Screen and Re-Screen; analysis of entire set of candidates:

*Muhammad Salim Hakeemi, Salim Ansari, Matthias Teuscher, Matthias Weißkopf*

Data handling and processing:

*Jürgen Dönitz*

Data collection and analysis for Drosophila comparison:

*Janna Siemanowski, Daniela Großmann, Muhammad Salim Hakeemi*

Annotation as part of last screening part:

*Xuebin Wan*

Generation and analysis of *Drosophila* mutant:

*Dorothea Schultheis*

Supervision and interpretation of Re-Screen:

*Martin Klingler, Michael Schoppmeier, Gregor Bucher*

*Siegfried Roth, Manfred Frasch*

Supervision primary Screen:

Coordination of screen funding and realization:

Martin Klingler, Michael Schoppmeier, Gregor Bucher

## Acknowledgements

We thank Mohamad Al Heshan, Elke Küster and Claudia Hinners for help with injection and processing. This project was funded by Deutsche Forschungsgemeinschaft (DFG research unit; FOR1234 iBeetle) and Bayer CropScience. The China Scholarship Council funded Xuebein Wan (201706760058).

## References

Ansari, S., Troelenberg, N., Dao, V. A., Richter, T., Bucher, G. and Klingler, M. (2018). Double abdomen in a short-germ insect: Zygotic control of axis formation revealed in the beetle Tribolium castaneum. PNAS 201716512.

Averof, M. (2002). Arthropod Hox genes: insights on the evolutionary forces that shape gene functions. Curr. Opin. Genet. Dev. 12, 386–392.

Bassett, A. R., Tibbit, C., Ponting, C. P. and Liu, J.-L. (2013). Highly efficient targeted mutagenesis of Drosophila with the CRISPR/Cas9 system. Cell Rep 4, 220–228.

Berghammer, A. J., Klingler, M. and Wimmer, E. A. (1999). A universal marker for transgenic insects. Nature 402, 370–1.

Brown, S., Fellers, J., Shippy, T., Denell, R., Stauber, M. and Schmidt-Ott, U. (2001). A strategy for mapping bicoid on the phylogenetic tree. Curr Biol 11, R43–4.

Bucher, G., Scholten, J. and Klingler, M. (2002). Parental RNAi in Tribolium (Coleoptera). Current Biology 12, R85–R86.

Carbon, S., Ireland, A., Mungall, C. J., Shu, S., Marshall, B., Lewis, S., the AmiGO Hub and the Web Presence Working Group (2009). AmiGO: online access to ontology and annotation data. Bioinformatics 25, 288–289.

Dönitz, J., Grossmann, D., Schild, I., Schmitt-Engel, C., Bradler, S., Prpic, N.-M. and Bucher, G. (2013). TrOn: An Anatomical Ontology for the Beetle Tribolium castaneum. PLOS ONE 8, e70695.

Dönitz, J., Schmitt-Engel, C., Grossmann, D., Gerischer, L., Tech, M., Schoppmeier, M., Klingler, M. and Bucher, G. (2015). iBeetle-Base: a database for RNAi phenotypes in the red flour beetle Tribolium castaneum. Nucl. Acids Res. 43, D720–D725.

Dönitz, J., Gerischer, L., Hahnke, S., Pfeiffer, S. and Bucher, G. (2018). Expanded and updated data and a query pipeline for iBeetle-Base. Nucleic Acids Res. 46, D831–D835.

Farnworth, M. S., Eckermann, K. N. and Bucher, G. (2019). Sequence heterochrony led to a gain of functionality in an immature stage of the central complex: a fly-beetle insight. bioRxiv 2019.12.20.883900.

Fu, J., Posnien, N., Bolognesi, R., Fischer, T. D., Rayl, P., Oberhofer, G., Kitzmann, P., Brown, S. J. and Bucher, G. (2012). Asymmetrically expressed axin required for anterior development in Tribolium. Proc. Natl. Acad. Sci. U.S.A. 109, 7782–7786.

Gilles, A. F., Schinko, J. B. and Averof, M. (2015). Efficient CRISPR-mediated gene targeting and transgene replacement in the beetle Tribolium castaneum. Development 142, 2832–2839.

Glinka, A., Wu, W., Delius, H., Monaghan, A. P., Blumenstock, C. and Niehrs, C. (1998). Dickkopf-1 is a member of a new family of secreted proteins and functions in head induction. Nature 391, 357–62.

Gurley, K. A., Rink, J. C. and Sánchez Alvarado, A. (2008). Beta-catenin defines head versus tail identity during planarian regeneration and homeostasis. Science 319, 323–327.

Herndon, N., Shelton, J., Gerischer, L., Ioannidis, P., Ninova, M., Dönitz, J., Waterhouse, R. M., Liang, C., Damm, C., Siemanowski, J., et al. (2020). Enhanced genome assembly and a new official gene set for Tribolium castaneum. BMC Genomics 21, 47.

Jorgensen, E. M. and Mango, S. E. (2002). The art and design of genetic screens: caenorhabditis elegans. Nat Rev Genet 3, 356–69.

Kile, B. T. and Hilton, D. J. (2005). The art and design of genetic screens: mouse. Nat. Rev. Genet. 6, 557–567.

Kittelmann, S., Ulrich, J., Posnien, N. and Bucher, G. (2013). Changes in anterior head patterning underlie the evolution of long germ embryogenesis. Dev. Biol. 374, 174–184.

Kitzmann, P., Schwirz, J., Schmitt-Engel, C. and Bucher, G. (2013). RNAi phenotypes are influenced by the genetic background of the injected strain. BMC Genomics 14, 5.

Kitzmann, P., Weißkopf, M., Schacht, M. I. and Bucher, G. (2017). A key role forfoxQ2in anterior head and central brain patterning in insects. Development 144, 2969–2981.

Klomp, J., Athy, D., Kwan, C. W., Bloch, N. I., Sandmann, T., Lemke, S. and Schmidt-Ott, U. (2015). Embryo development. A cysteine-clamp gene drives embryo polarity in the midge Chironomus. Science (New York, N.Y.) 348, 1040–1042.

Kondo, S. and Ueda, R. (2013). Highly improved gene targeting by germline-specific Cas9 expression in Drosophila. Genetics 195, 715–721.

Lorenzen, M. D., Kimzey, T., Shippy, T. D., Brown, S. J., Denell, R. E. and Beeman, R. W. (2007). piggyBac-based insertional mutagenesis in Tribolium castaneum using donor/helper hybrids. Insect Mol Biol 16, 265–275.

Lynch, J. A. and Roth, S. (2011). The evolution of dorsal–ventral patterning mechanisms in insects. Genes Dev. 25, 107–118.

Mungall, C. J., Gkoutos, G. V., Smith, C. L., Haendel, M. A., Lewis, S. E. and Ashburner, M. (2010). Integrating phenotype ontologies across multiple species. Genome Biol 11, R2.

Panfilio, K. A. (2008). Extraembryonic development in insects and the acrobatics of blastokinesis. Developmental Biology 313, 471–491.

Patton, E. E. and Zon, L. I. (2001). The art and design of genetic screens: zebrafish. Nat Rev Genet 2, 956–66.

Posnien, N., Schinko, J. B., Kittelmann, S. and Bucher, G. (2010). Genetics, development and composition of the insect head - A beetle’s view. Arthropod Struct Dev 39, 399–410.

Schinko, J. B., Kreuzer, N., Offen, N., Posnien, N., Wimmer, E. A. and Bucher, G. (2008). Divergent functions of orthodenticle, empty spiracles and buttonhead in early head patterning of the beetle Tribolium castaneum (Coleoptera). Dev Biol 317, 600–13.

Schinko, J. B., Weber, M., Viktorinova, I., Kiupakis, A., Averof, M., Klingler, M., Wimmer, E. A. and Bucher, G. (2010). Functionality of the GAL4/UAS system in Tribolium requires the use of endogenous core promoters. BMC Dev Biol 10, 53.

Schmitt-Engel, C., Schultheis, D., Schwirz, J., Strohlein, N., Troelenberg, N., Majumdar, U., Dao, V. A., Grossmann, D., Richter, T., Tech, M., et al. (2015). The iBeetle large-scale RNAi screen reveals gene functions for insect development and physiology. Nat Commun 6,.

Schultheis, D. (2016). Identifizierung und Charakterisierung neuer regulatorischer Gene in der Muskelentwicklung durch einen genomweiten RNAi-Screen in Tribolium castaneum.

Schultheis, D., Weißkopf, M., Schaub, C., Ansari, S., Dao, V. A., Grossmann, D., Majumdar, U., Hakeemi, M. S., Troelenberg, N., Richter, T., et al. (2019a). A Large Scale Systemic RNAi Screen in the Red Flour Beetle Tribolium castaneum Identifies Novel Genes Involved in Insect Muscle Development. G3 (Bethesda) 9, 1009–1026.

Schultheis, D., Schwirz, J. and Frasch, M. (2019b). RNAi Screen in Tribolium Reveals Involvement of F-BAR Proteins in Myoblast Fusion and Visceral Muscle Morphogenesis in Insects. G3 (Bethesda) 9, 1141–1151.

Snodgrass, R. (1954). Insect Metamorphosis: Smithsonian Miscellaneous Collections, V122, No. 9. Washington: Literary Licensing.

St Johnston, D. (2002). The art and design of genetic screens: Drosophila melanogaster. Nat Rev Genet 3, 176–88.

Stappert, D., Frey, N., von Levetzow, C. and Roth, S. (2016). Genome-wide identification of Tribolium dorsoventral patterning genes. Development 143, 2443–2454.

Tautz, D., Friedrich, M. and Schröder, R. (1994). Insect embryogenesis – what is ancestral and what is derived? Development 1994, 193–199.

Tomoyasu, Y. and Denell, R. E. (2004). Larval RNAi in Tribolium (Coleoptera) for analyzing adult development. Dev Genes Evol 214, 575–8.

Weigel, D., Jürgens, G., Kuttner, F., Seifert, E. and Jäckle, H. (1989). The homeotic gene fork head encodes a nuclear protein and is expressed in the terminal regions of the Drosophila embryo. Cell 57, 645–58.

Wimmer, E. A., Cohen, S. M., Jackle, H. and Desplan, C. (1997). buttonhead does not contribute to a combinatorial code proposed for Drosophila head development. Development 124, 1509–17.

Yoon, Y., Klomp, J., Martin-Martin, I., Criscione, F., Calvo, E., Ribeiro, J. and Schmidt-Ott, U. (2019). Embryo polarity in moth flies and mosquitoes relies on distinct old genes with localized transcript isoforms. Elife 8,.

