## Supplementary Figures 1-3 for "Large portion of essential genes is missed by screening either fly or beetle indicating unexpected diversity of insect gene function"

### Figure S1: Genes known from Drosophila tested in the iBeetle screen

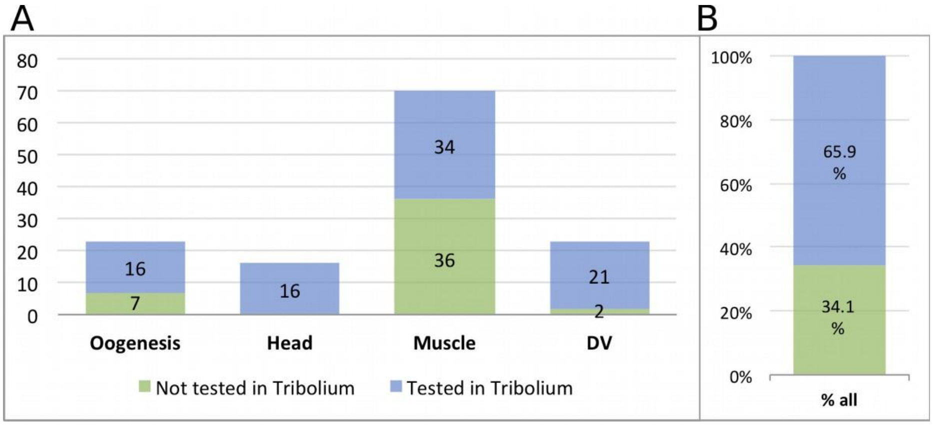


### Figure S2: Phylogenetic_Tree_TC001720

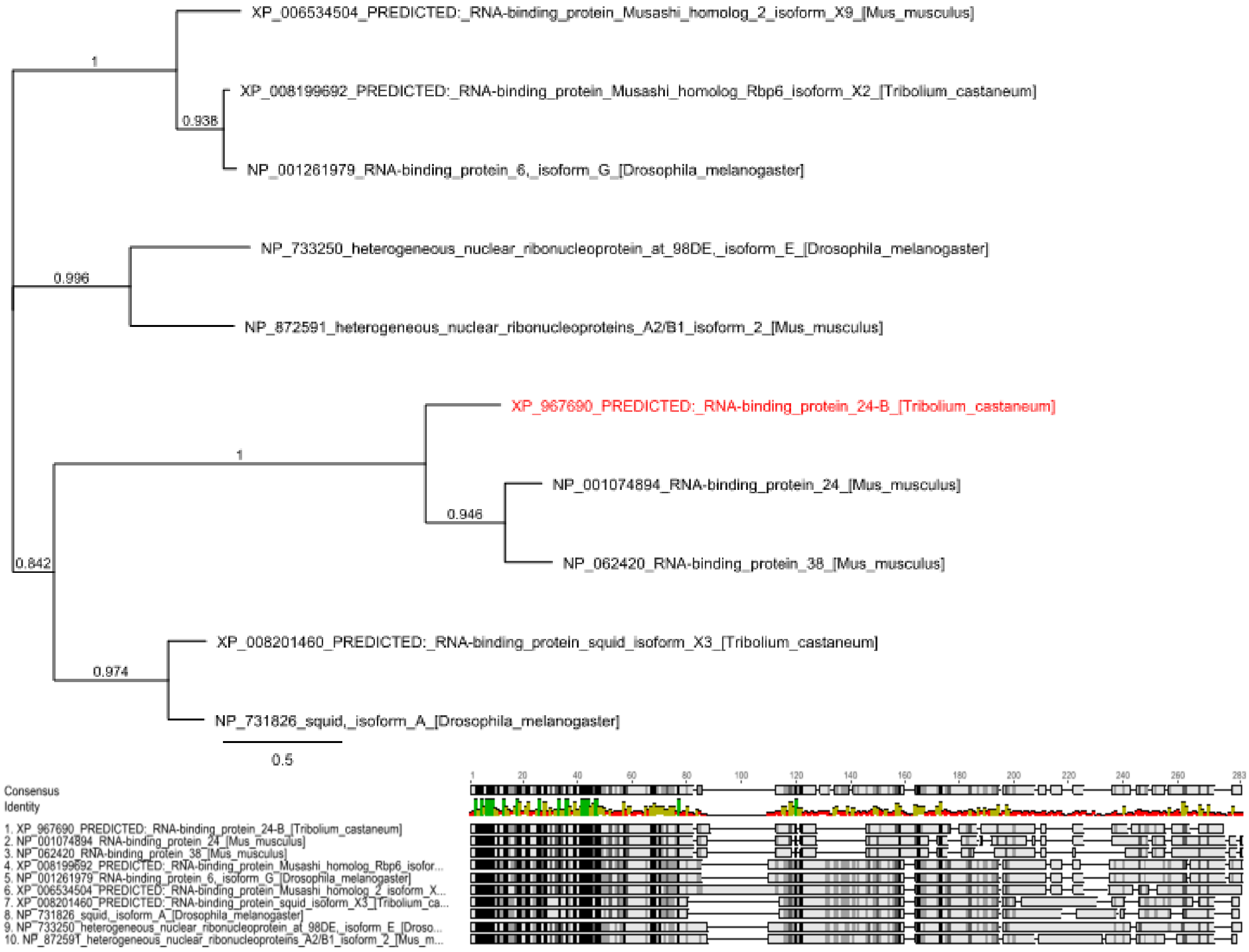

### Figure S3: Phylogenetic_Tree_TC002909

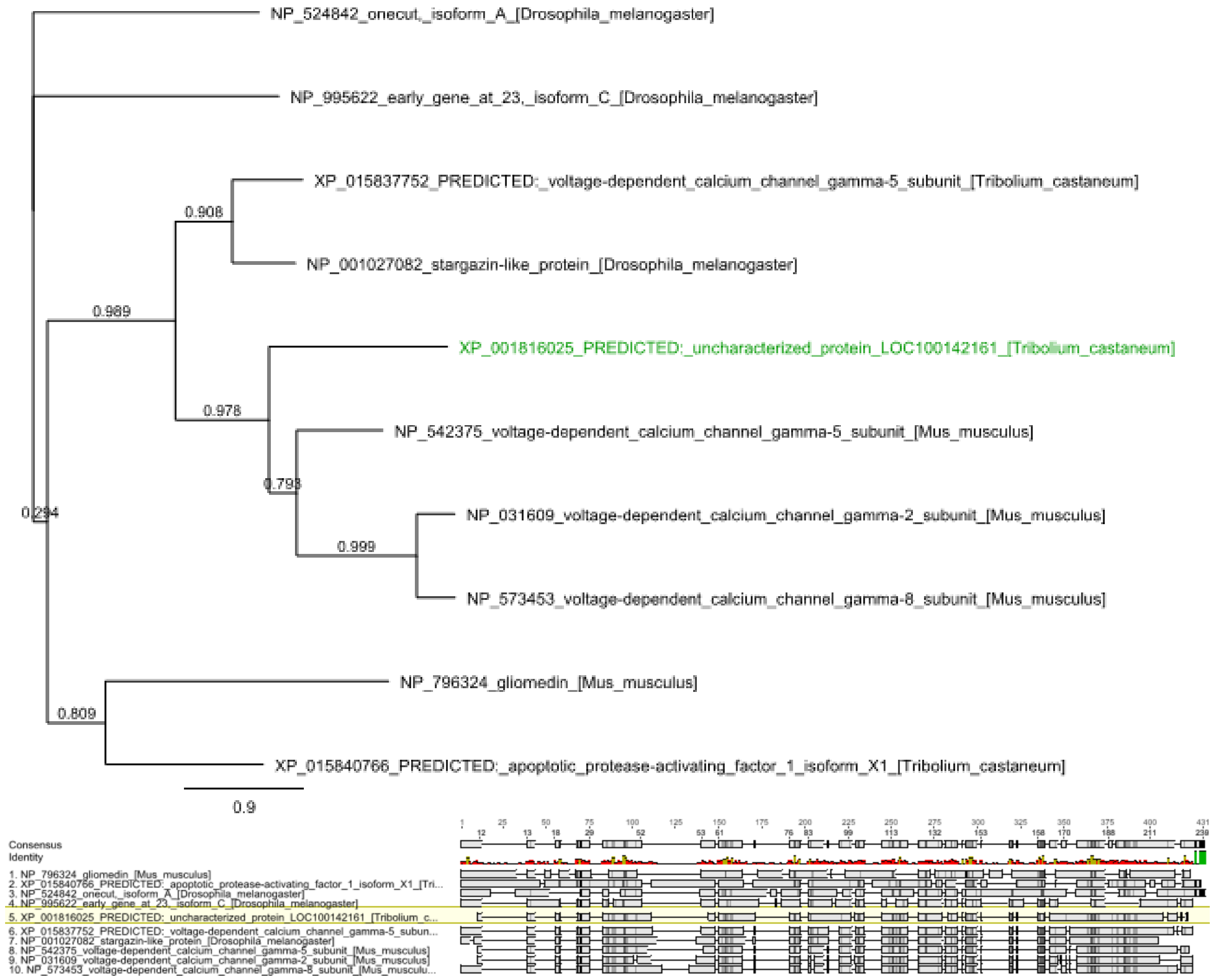
